# Differentiating Genomic SNPs Using Allele Depth and Predicted Genotype

**DOI:** 10.1101/438325

**Authors:** Saam Hasan

**Author notes:** Corresponding author: Saam Hasan, Mailing address: Plot 15, Block B, Bashundhara, Dhaka 1229, Bangladesh.

## Abstract

Differentiating between genomic SNPs and other types of single nucleotide variants becomes a key issue in research aimed at studying the importance of these variants of a particular type in biological processes. Here we present an R based method for differentiating between genomic single nucleotide polymorphisms (SNPs) and RNA editing sites. We use data from an earlier study of ours and target only the known dbsnp SNPs that we found in our study. Our method involves calculating the ratio of allele depth for ref and alt alleles and comparing that to the predicted genotype. We use the concept that editing levels should be different for each allele and thus should not reflect the ratio predicted by the genotype. The study yielded an accuracy rate ranging from 86 to over 90 percent at successfully predicted dbsnp entries as SNPs. Albeit this is in the absence of known RNA editing site vcf data to compare as a reference.

## Introduction

Computational techniques for the analysis of next generation sequencing data has revolutionized the way biological data can be interpreted and used. Techniques have been developed for all kinds of uses with all categories of biological data such as genome sequences, transcriptome sequences, metagenomics, proteomics, metabolomics, etc. RNA data has occupied a special significance to research interests due to RNA’s central role in gene expression and protein translation as well as a host of regulatory purposes such as micro RNAs and other processes such as RNA editing (2). The development of tools for the analysis of RNA sequence data has been rapid and led to the availability of a host of toolkits for analysis, validation and interpretation of RNA sequences. However one major challenge has been to differentiate sequence changes in RNA that occurred post transcriptionally from those occurred at the genomic level. In most cases both DNA and RNA sequences for the subject of the study are used to differentiate such changes. Although tools have been developed that can do so from RNA sequence data alone.

Here we define an R based method for differentiating between genomic single nucleotide variants (SNPs) from other kinds of changes (most RNA editing sites). Our method relies on information about genotype and allele count available in a variant call format (vcf) file which is the most common and accepted means of representing single nucleotide variant data (3). The tool is designed to work with vcf version 4.2 but should work with other versions as well subject to a few tweaks to the code. It uses the predicted genotype in the vcf and compares it to the actual number of alleles observed for the reference and alternate nucleotides at each variant call position. As explained by Zhang and Xiao the major difference between RNA editing sites and genomic SNPs are variable editing levels for the two alleles of each variant. The two alleles are most likely to be edited differentially and hence the ratio of observed alleles should not reflect the ratio predicted by the genotype (4). We use this method for heterozygous variants only. Homozygous variants are normally better accepted as non-Genomic SNPs (5). Here we apply this method to data from a study whose pre-print version we published earlier this year (6). Out of that data, we only applied this for the known genomic SNPs found in our variant calls and not our identified novel sites (as the latter remain unverified still). We predicted the nature of each variant in our vcf files and then focussed on the dbsnp (8) variants present there in order to ascertain the power of the method for correctly calling SNPs.

## Methods

Vcf files contain information regarding a predicted genotype at each variant call position (the GT field in the Genotype column). Variant callers such haplotypeCaller and UnifiedGenotyper from GATK use their algorithms to calculate the most likely genotype at each locus. The AD field then states the depth for each allele. The latter refers to the number of reads that harboured that particular variant (shown separately for ref and alt alleles) (3). Our R based method first parses the vcf file using the bedr package available on R (10). Then it uses the stringr package to split the genotype fields so that can be easily manipulated (9). Afterwards we create two separate fields for the ref alt read depth ratio and then the type of variant. The ref alt read depth ratio is simply obtained by dividing the read depth for ref by the read depth for alt. We then focus on the heterozygous variants only as the target of our study as previously explained. As explained by the Zhang and Xiao, this ratio should not agree with the predicted genotype (4). For heterozygous calls (0/1), we selected a cut off 2 and 0.5 beyond which we considered the ratios as deviating from the genotype. The former for when the ref allele read depth > alt allele read depth and the latter for when the condition is reversed. For homozygous calls, we simply took the presence of any ref alleles as deviation from the ratio predicted by genotype (i.e. ref alt read depth ratio > 0). Reads with alt allele depth 0 (uninformative reads) were discarded from consideration. Finally, homozygous variants with ref alt ratio greater than zero were counted as RNA editing sites and the rest as SNPs. Heterozygous variants with ref alt ratio greater than 2 (for ref read depth > alt read depth) or less than 0.5 (for ref read depth < alt read depth) were counted as editing sites and the rest as SNPs. We applied this method to seven datasets from a previous research whose pre-print version we published earlier this year. As the new variants identified are not yet verified, we focussed only on the known variants in these datasets and calculated the ability of this method to correctly call them as SNPs. The five datasets were from a study conducted by *Labonte et al* in 2017 on the effect of sex specific transcriptional regulators in Major Depressive Disorder (MDD) (7). In order to identify potential RNA editing sites unique to the individuals with MDD, we had applied a variant calling pipeline to seven of these datasets (6).

## Results

Table 1 lists the total number of dbsnp variants present in the datasets and how many of them were called correctly by our method. The percentage of correct calls for dbsnp entries as SNPs ranged from 86 to over 90%. For each of the datasets labelled Runs 1, 3, 4, 5, 6, 12 and 13, the percentage of dbsnp variants correctly called as SNPs were 86.4, 88.9, 90.1, 86.2, 87.7, 87.7, 86.9 respectively. Figure 1a to 1g shows the proportion of dbsnp entries that were correctly detected. In the figures, undetected refers to the small outer portion of the Venn diagram and the number states how many dbsnp entries were undetected as SNPs. The majority of non dbsnp entries were also called as SNPs. The remaining variant calls were either uninformative or called as editing sites based on the rationale described in methods. Most of the SNP calls were homozygous variants with only the alt allele being observed.

**Table 1:** Percentage of dbnsps in each dataset called correctly.

| Run | Run1 | Run3 | Run4 | Run5 | Run6 | Run12 | Run13 |
| --- | --- | --- | --- | --- | --- | --- | --- |
| Total | 606938 | 401223 | 270214 | 599935 | 530667 | 545738 | 703541 |
| Total Without dnsp | 374468 | 214035 | 137447 | 329086 | 293650 | 295502 | 382241 |
| Dbsnp only | 232470 | 187188 | 132767 | 270849 | 237017 | 250236 | 321300 |
| Total Dbsnp correctly called | 200924 | 166531 | 119635 | 233645 | 207870 | 219656 | 279271 |

**Figure 1a:**
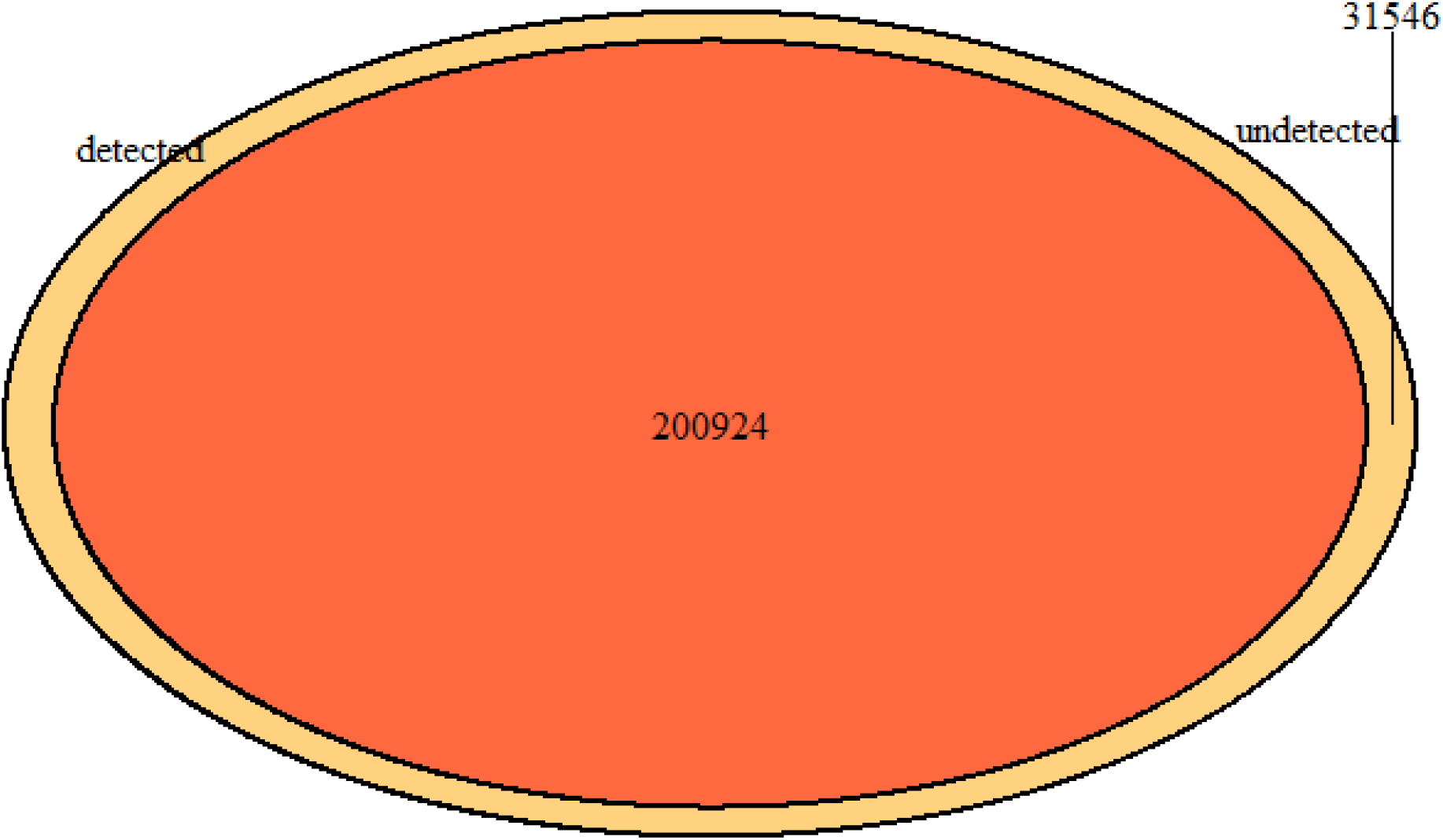
Correct and incorrect calls for Run1.

**Figure 1b:**
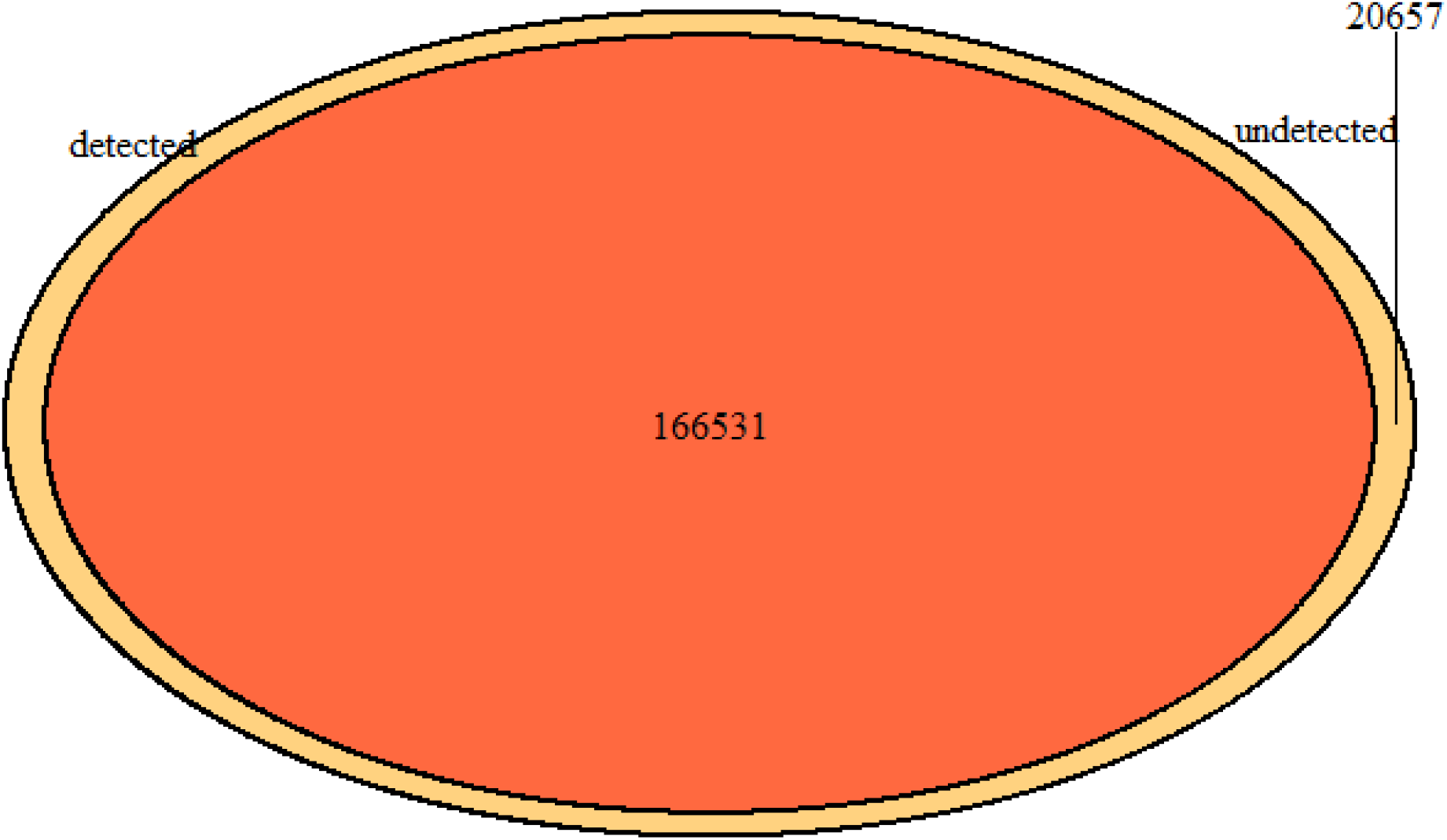
Correct and incorrect calls for Run3.

**Figure 1c:**
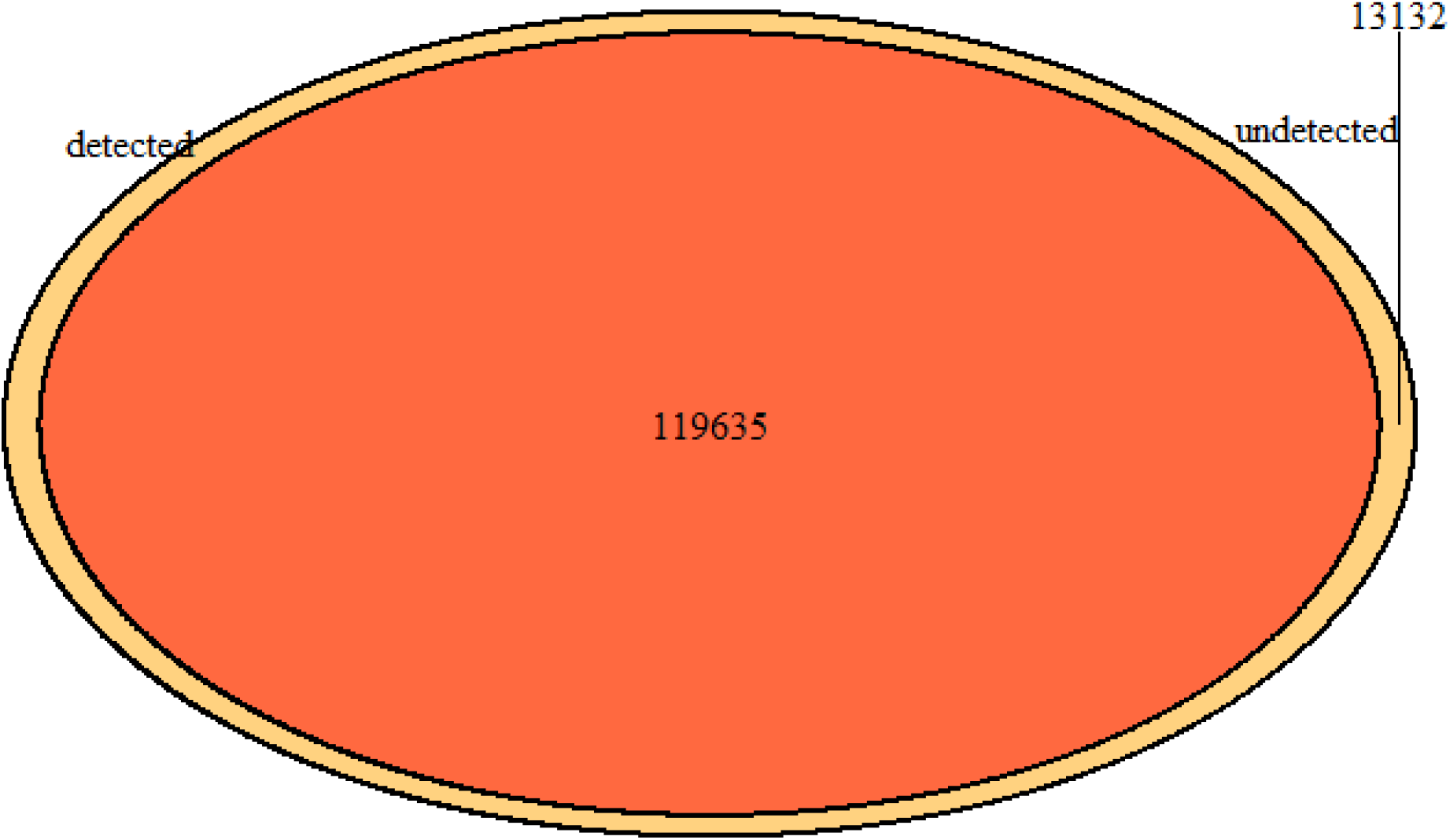
Correct and incorrect calls for Run4

**Figure 1d:**
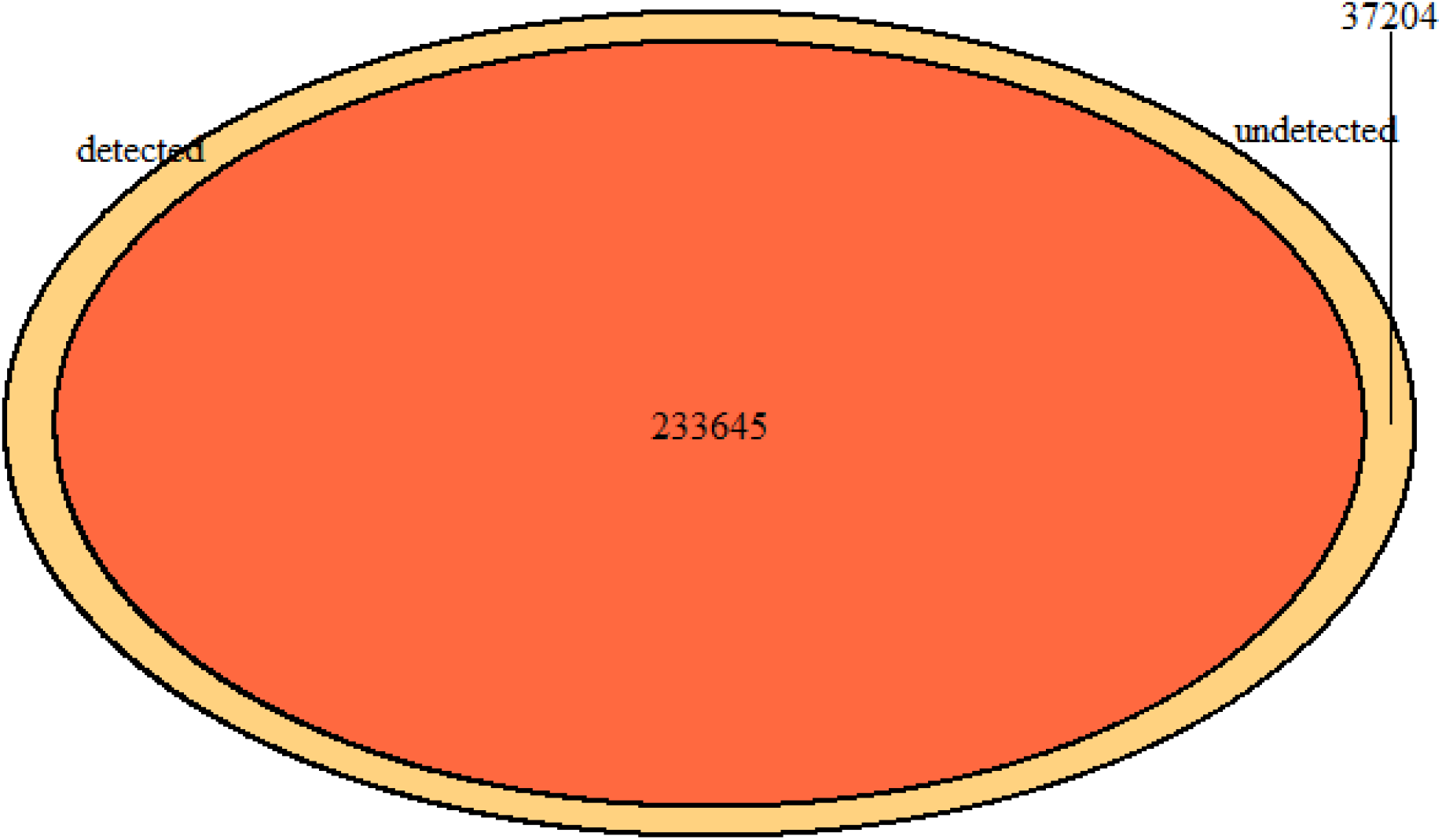
Correct and incorrect calls for Run5

**Figure 1e:**
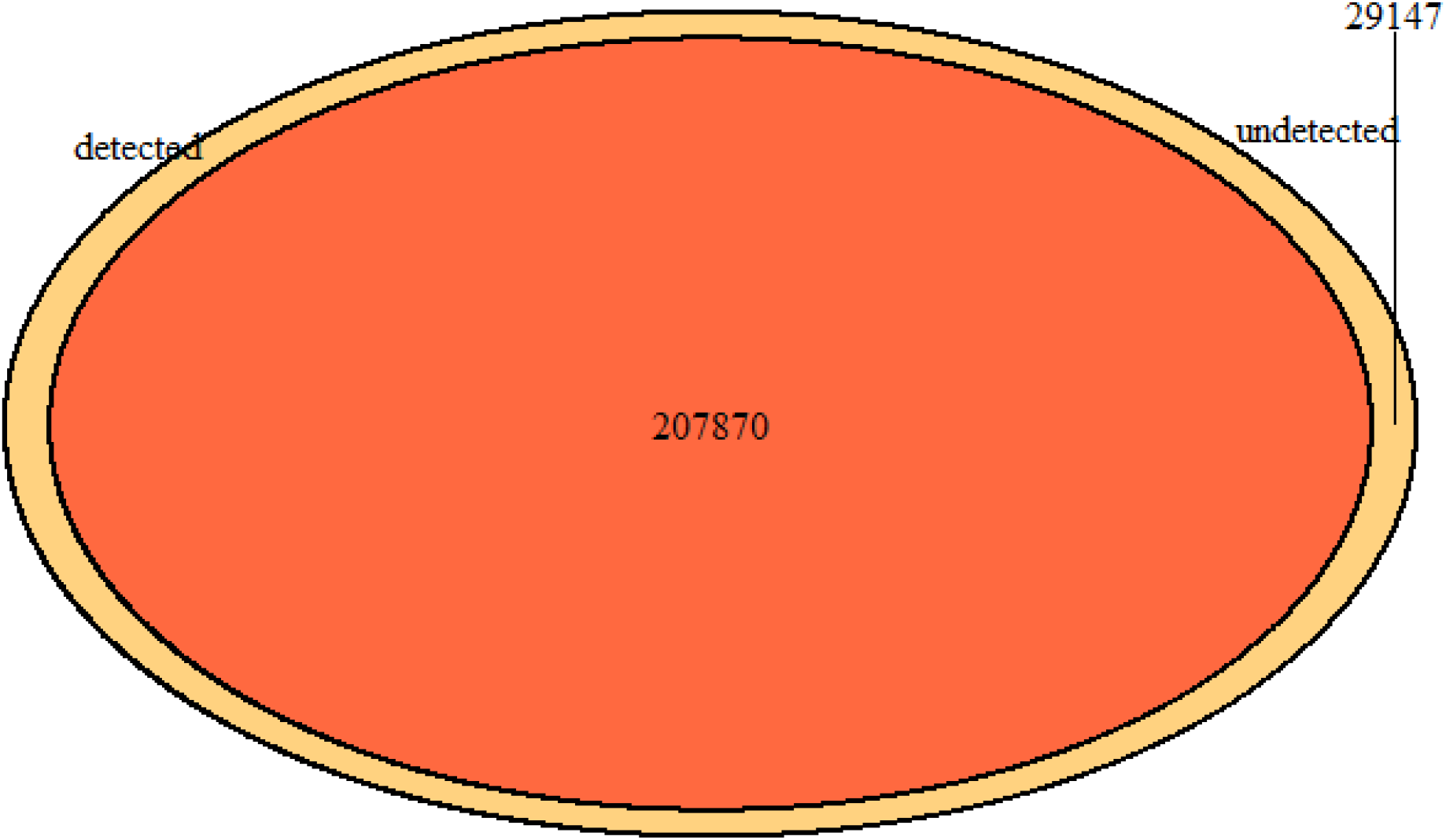
Correct and incorrect calls for Run6

**Figure 12:**
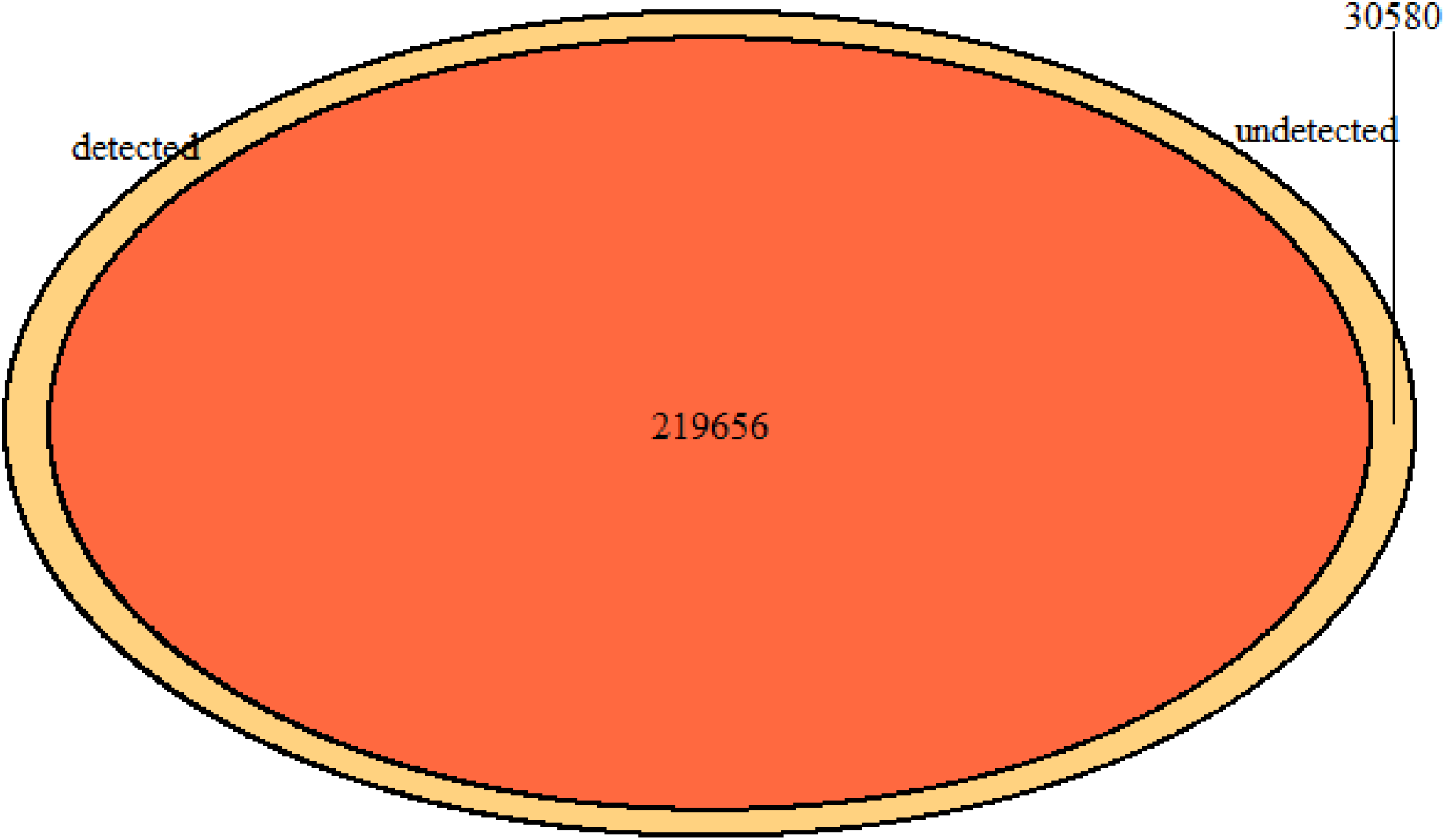
Correct and incorrect calls for Run12

**Figure 1g:**
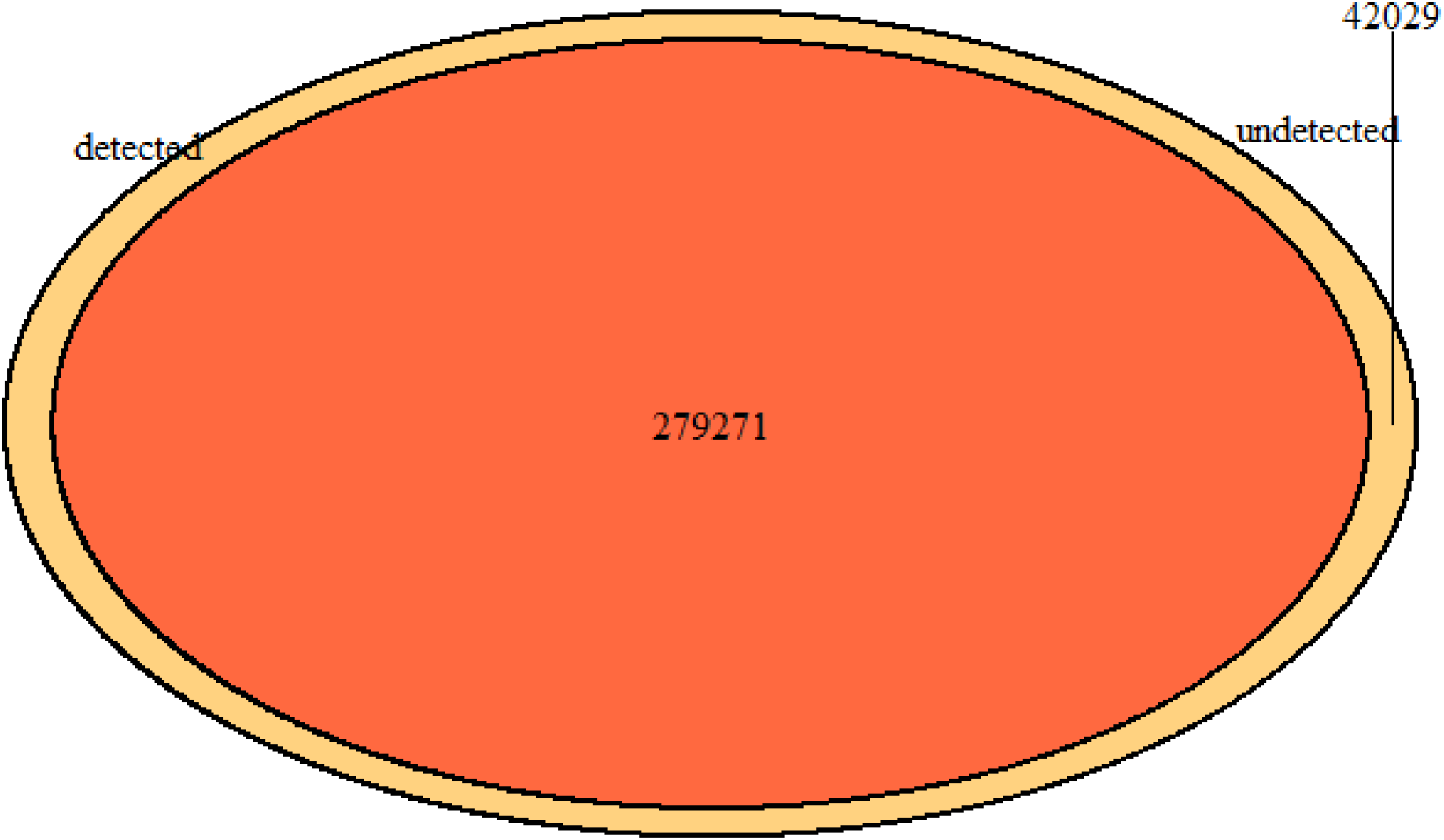
Correct and incorrect calls for Run13

## Discussion

Discriminating between types of single nucleotide variants often represents a key issue when working with such data while investigating a particular type of variant. Here we described a method for doing so using allele depth and predicted genotype. Although this is still at a rudimentary stage and not fully proven in the absence of a reference RNA editing site vcf to test on, the preliminary results at predicting dbsnps are positive. The majority of variants in the datasets were homozygous and thus not taken into consideration for our analysis. In addition to this, other information may also be taken into consideration for discriminating between variants. We compared one of our datasets (run 4) with those known RNA editing sites and did obtain a handful of common sites (25 in total). Although in too small a number for any meaningful deduction to be made, we did find a difference in the BaseQRankSum parameter between the editing sites and most dbsnps. This parameter which is the allele specific Z score value from Wilcoxon rank sum test of each Alt Vs. Ref base qualities, was failed by a large number of the dbsnps while most of our very small number of known editing sites passed it. The availability of large RNA editing site datasets in vcf format would enable this and other possible differentiating features to be evaluated. This method may have applications in research where either type of variant is desired. In particular if its efficiency at differentiating genomic SNPs from other types of RNA or post transcriptional variants can be established, this can be used in research where either genomic SNPs or the RNA derived variants are the object of study.

## Availabiliy

The script and a sample dataset are available at this github repository: https://github.com/SaamH94/Variant-Predictor

## References

1. Wadapurkar R, Vyas R. Computational analysis of next generation sequencing data and its applications in clinical oncology. Informatics in Medicine Unlocked. 2018;11:75–82.

2. Maydanovych O, Beal P. Breaking the Central Dogma by RNA Editing. ChemInform. 2006;37(44).

3. DePristo M, Banks E, Poplin R, Garimella K, Maguire J, Hartl C et al. A framework for variation discovery and genotyping using next-generation DNA sequencing data. Nature Genetics. 2011;43(5):491–498.

4. Zhang Q, Xiao X. Genome sequence–independent identification of RNA editing sites. Nature Methods. 2015;12(4):347–350.

5. Ramaswami G, Lin W, Piskol R, Tan M, Davis C, Li J. Accurate identification of human Alu and non-Alu RNA editing sites. Nature Methods. 2012;9(6):579–581.

6. Hasan S, Robbani S, Afroze T, Ahsan G, Hossain M. Identifying Changes in RNA Editome Unique to Major Depressive Disorder. 2018;.

7. Labonté B, Engmann O, Purushothaman I, Menard C, Wang J, Tan C et al. Sex-specific transcriptional signatures in human depression. Nature Medicine. 2017;23(9):1102–1111.

8. Smigielski E. dbSNP: a database of single nucleotide polymorphisms. Nucleic Acids Research. 2000;28(1):352–355.

9. Wickham H (2018). stringr: Simple, Consistent Wrappers for Common String Operations. R package version 1.3.1. htps://CRAN.R-project.org/package=stringr

10. Waggott D, Haider S, Paul C. Boutros(2017). bedr: Genomic Region Processing using Tools Such as ‘BEDTools’, ‘BEDOPS’, ‘TABIX’. R package version 1.0.4. https://CRAN.R-project.org/package=bedr.

